# Equilibrium dynamics of a biomolecular complex analyzed at single-amino acid resolution by cryo-electron microscopy

**DOI:** 10.1101/2022.06.24.497460

**Authors:** Daniel Luque, Alvaro Ortega-Esteban, Alejandro Valbuena, Jose Luis Vilas, Alicia Rodríguez-Huete, Mauricio G. Mateu, José R. Castón

## Abstract

The biological function of macromolecular complexes depends not only on large-scale transitions between conformations, but also on small-scale conformational fluctuations at equilibrium. This study shows that detailed information on the equilibrium dynamics of a biocomplex obtained from local resolution (LR) data in cryo-electron microscopy (cryo-EM) density maps matches the information obtained in solution by hydrogen-deuterium exchange mass-spectrometry. The structure of the minute virus of mice (MVM) capsid as a model system was determined by cryo-EM, and LR values were found to correlate with both crystallographic B-factors and hydrogen-deuterium exchange values. Once validated, the cryo-EM LR-based approach was used to compare the equilibrium dynamics of wild-type and two mutant MVM capsids. The results revealed a linkage between mechanical stiffening and impaired equilibrium dynamics in a biocomplex. Cryo-EM is emerging as a powerful approach for simultaneously acquiring information on the atomic structure and local equilibrium dynamics of biomolecular complexes.

## Introduction

Macromolecular complexes carry out many biochemical processes in cells and organisms. These nanomachines convert energy into atomic movements, which are generally reflected in conformational changes that mediate biological functions. Viruses and their capsids have emerged as a paradigm to study the conformational dynamics of biomolecular complexes and its biological relevance (Johnson, 2003; Sherman, Smith, & Smith, 2020).

During virus morphogenesis, multiple copies of one or a few capsid proteins (CP) self-assemble into a protein shell (capsid) that encloses the viral genome to yield an infectious virus particle (Harrison, 2007; Luque & Caston, 2020; Mateu, 2013). The built-in conformational flexibility of CPs and capsids is required to fulfill their specific roles in the viral infectious cycle (Bothner & Hilmer, 2011; Johnson, 2003). Most studies on virus dynamics have focused in large-scale transitions between populated states of the viral particle. For example, during virus maturation a labile capsid may undergo a large, irreversible structural transition to become a robust but metastable capsid. The mature capsid effectively protects the viral genome in the extracellular medium, but allows its release in the host cell (Veesler & Johnson, 2012).

In addition to transitions between different thermodynamic states, viral capsids are subjected to conformational fluctuations at equilibrium, termed equilibrium dynamics, that are also relevant for viral infection (Bothner & Hilmer, 2011). These fluctuations involve continuous changes in the position of atoms, atomic groups and larger structural elements. For example, capsid “breathing” allows the externalization of certain CP regions that carry signals required for infection by some viruses (Bothner, Dong, Bibbs, Johnson, & Siuzdak, 1998; Bothner & Hilmer, 2011; Bothner et al., 1999; Lewis, Bothner, Smith, & Siuzdak, 1998; Roivainen, Piirainen, Rysa, Narvanen, & Hovi, 1993). Conformational fluctuation at equilibrium is a biologically critical feature also of many other biomolecular machines (Agarwal, Bernard, Bafna, & Doucet, 2020; DuBay, Bowman, & Geissler, 2015; Hodge, Benhaim, & Lee, 2020; Ross et al., 2022).

Different approaches have been used to investigate large-scale conformational transitions during virus assembly or maturation, or during viral genome packaging or uncoating. Stable populated intermediates and kinetically trapped intermediates have been structurally characterized (Bothner & Hilmer, 2011), sometimes using X-ray crystallography or cryo-electron microscopy (cryo-EM) to reach atomic resolution (Veesler & Johnson, 2012; C. Wang, Zeng, & Wang, 2022; Wikoff et al., 2000; Woodson et al., 2021; Xu et al., 2021). Fewer techniques are suited to probe the small-scale equilibrium dynamics of viruses and other large molecular complexes. Binding of specific antibodies has been used to infer capsid breathing from the transient externalization of internally located CP segments (Bothner & Hilmer, 2011; Roivainen et al., 1993). Covalent crosslinking of specific capsid residues, limited proteolysis or hydrogen-deuterium exchange (HDX) combined with mass spectrometry (MS) have identified highly dynamic structural elements in some viral capsids (Bothner et al., 1998; Bothner & Hilmer, 2011; Bothner et al., 1999; Lewis et al., 1998; van de Waterbeemd et al., 2017; L. Wang & Smith, 2005; Worner, Shamorkina, Snijder, & Heck, 2021). These approaches have provided many important insights into the equilibrium dynamics of viruses and other biocomplexes. However, they may be technically intricate, may probe only some capsid regions, and do not reach atomic resolution.

Nuclear magnetic resonance (NMR) spectroscopy has also been used to study the dynamics of virus particles (Szymczyna, Gan, Johnson, & Williamson, 2007). NMR can reach near-atomic resolution when probing the dynamics of smaller proteins or some specially arranged supramolecular assemblies such as filamentous phages (Opella, Zeri, & Park, 2008). However, technical limitations do not generally allow NMR spectroscopy to provide a complete, near-atomic resolution description of the equilibrium dynamics of most viruses and macromolecular assemblies. All-atom molecular dynamics (MD) simulations have predicted at atomic resolution the fast equilibrium dynamics of some virus particles (Freddolino, Arkhipov, Larson, McPherson, & Schulten, 2006), but severe computational demands, inaccuracies in the force field and other issues currently impose substantial limitations to MD for studying the equilibrium dynamics of viruses and other large molecular complexes.

Cryo-EM is greatly contributing to our understanding of viruses and cellular machines by providing outstanding structural descriptions at atomic or near-atomic resolution (Kaelber, Hryc, & Chiu, 2017; Zhou, 2011). However, as for X-ray crystal structures, cryo-EM-derived structures generally correspond to either the minimum free energy conformation (under the conditions used), or to a kinetically trapped state. Thus, it has been argued that these two techniques provide only static, somewhat deceptive views of macromolecular complexes. In fact, both techniques can provide also relevant, high-resolution information on their conformational dynamics.

For each atom in a crystallized molecule a B-factor (or “temperature factor”) can be calculated from diffraction data. If interpreted with caution, a set of relative (normalized) B-factors for a molecular complex may provide an approximate description at near-atomic resolution of spatial differences in small-scale conformational fluctuations at equilibrium (Sun, Liu, Qu, Feng, & Reetz, 2019). Likewise, a few approaches have recently been developed to extract, from cryo-EM data, information on the conformational dynamics of viruses and other biocomplexes. For example, rapid vitrification may trap some individual particles of a macromolecular complex in transient, higher free energy conformations (Dandey et al., 2020; Frank, 2017). If these conformations are sufficiently different from the minimum free energy conformation and from each other, structural models for different, inter-converting conformations may be obtained from the corresponding electron density maps, and compared to identify highly dynamic regions. In addition, sophisticated computational approaches have been applied to investigate dynamics from a continuous distribution of heterogeneities in data from an ensemble of individual cryo-EM images (Bonomi, Pellarin, & Vendruscolo, 2018; Doerschuk, Gong, Xu, Domitrovic, & Johnson, 2016; Hartmann et al., 2021; Liao, Hashem, & Frank, 2015; Oide, Kato, Oroguchi, & Nakasako, 2020; Penczek, Kimmel, & Spahn, 2011; Tang et al., 2014; Q. Wang et al., 2013). However, the detailed information on equilibrium dynamics that can potentially be extracted from cryo-EM data had not been validated by comparison with equilibrium dynamics information determined in solution using hydrogen-deuterium exchange (HDX-MS), a gold-standard technique.

The present study validates, by direct comparison with HDX-MS results, a simple, general approach to extract, from local resolution differences (LR) in the cryo-EM density map, relevant information about the small-scale equilibrium dynamics of supramolecular complexes. For proof of concept, in the second part of this study, the validated cryo-EM approach has been applied to investigate a suggested linkage between changes in the mechanical elasticity of a viral capsid, and changes in its equilibrium dynamics leading to impaired biological function.

## Results

### Equilibrium dynamics of the minute virus of mice (MVM) capsid revealed by local resolution analysis of cryo-EM maps

The parvovirus MVM is one of the smallest (∼250 Å in diameter) and structurally simplest viruses known, yet it fulfills many complex functions leading to infection of host cells (Cotmore & Tattersall, 2014). Thus, this virus constitutes an excellent system to investigate general relationships between structure, dynamics and function of viruses and biomolecular complexes in general. The icosahedral T=1 MVM capsid is built from 60 CP subunits with identical sequence and tertiary structure, except for a N-terminal (Nt) extension that in 10 (VP1) subunits is longer than in the other 50 (VP2) subunits. The initially internal, structurally disordered VP1 extensions do not contribute to capsid structure or assembly. VP2-only MVM capsids made of 60 identical subunits (Hernando et al., 2000) have previously facilitated the study of different structure-function relationships, and were also used in this study. The atomic structure of the VP2-only capsid of the wild-type (wt) MVMp (prototype strain p) had previously been determined by X-ray crystallography (Kontou et al., 2005). In addition, the equilibrium dynamics in solution of nearly every peptide segment in exactly the same capsid had previously been analyzed by HDX-MS (van de Waterbeemd et al., 2017). The results of those studies made this viral capsid a model of choice to assess whether reliable, high-resolution information on equilibrium dynamics could be derived from LR data in a cryo-EM density map of a supramolecular complex.

#### Near-atomic-resolution cryo-EM structure of the MVM capsid

As a first step, the atomic structure of the wt MVMp VP2-only capsid was determined by cryo-EM. Highly purified MVM capsids were visualized in a 200-kV FEI Talos Arctica electron microscope (Fig. 1A). We merged ∼210,000 particle images to calculate an icosahedrally averaged map at 3.42 Å resolution (Fig. 1B), as estimated by the criterion of 0.143 Fourier shell correlation (FSC) coefficient (Fig. S1). Using the crystal structure of the same MVM capsid [PDB ID: 1Z14; (Kontou et al., 2005)] as initial template, the polypeptide chain was built using Coot (Emsley & Cowtan, 2004). The near-atomic model obtained for VP2 includes 549 residues (Gly39-Tyr587) of the total 587 residues; the 38 Nt residues are invisible due to extensive disorder.

**Figure 1.**
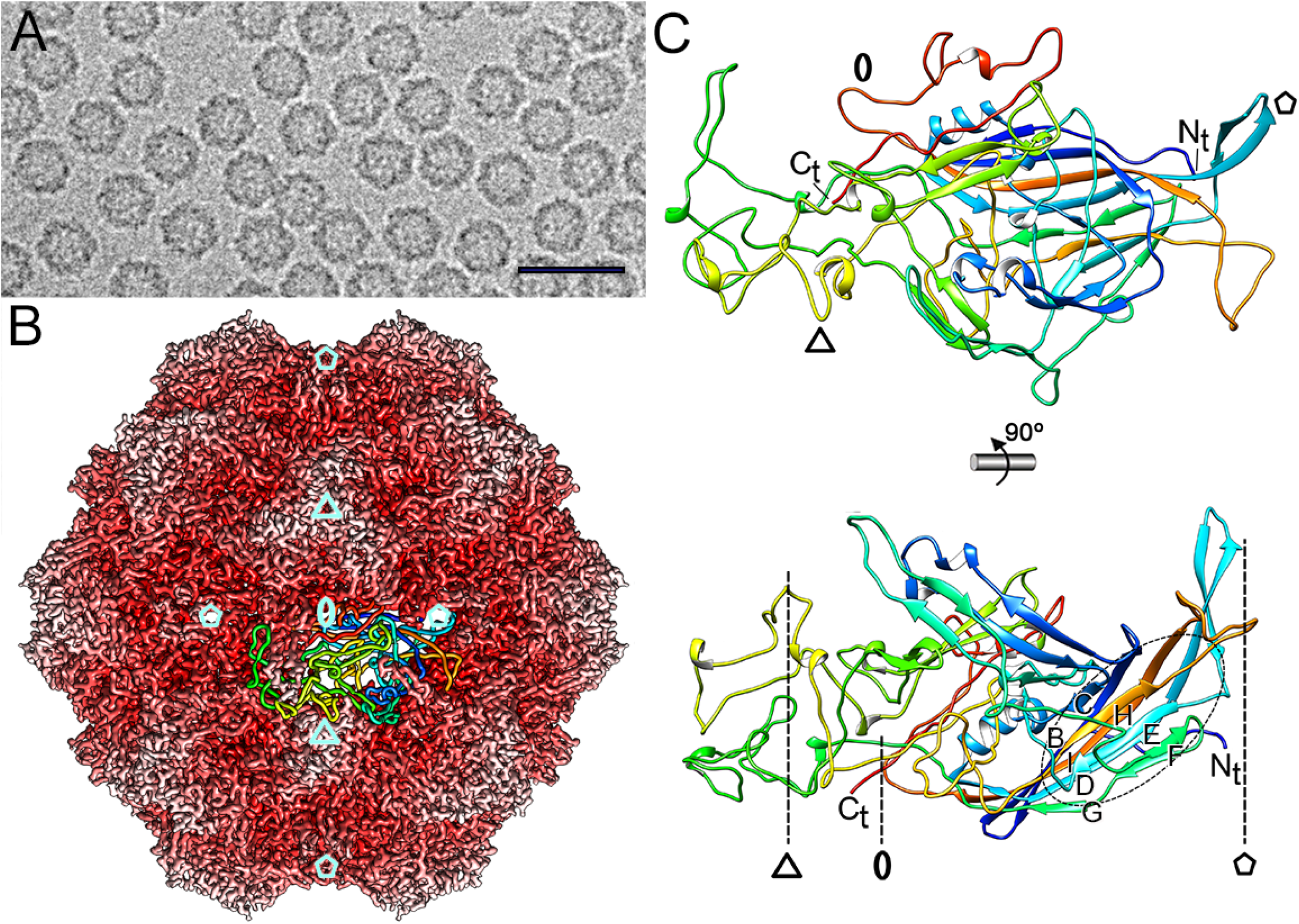
Cryo-EM structure of the wt MVM capsid. (A) Cryo-electron micrograph of wt MVM capsids. Bar, 50 nm. (B) Radially color-coded atomic model of the wt MVM capsid viewed along a 2-fold axis. Protruding trimers (white) are visible in the T=1 lattice. The atomic structure of a CP subunit is shown. Cyan pentagons, triangles or ovals respectively indicate 5-fold, 3-fold or 2-fold icosahedral symmetry axes. These axes define S5, S3 or S2 regions in the capsid, respectively. (C) Ribbon diagram of the CP monomer (top view, top; side view, bottom). The CP subunit is rainbow-colored from blue (N terminus [N_t_]) to red (C terminus [C_t_]). Dashed oval indicates the VP2 β-barrel. Symbols as defined for panel (A) indicate icosahedral symmetry axes.

In the cryo-EM model obtained the structural features of the CP (VP2) could be observed in atomic detail. The CP is folded as a single β-barrel (the jelly-roll fold) with two four-stranded β-sheets facing each other (βB to βI) tangential to the capsid surface (Fig. 1C, bottom). Loop insertions between β-strands are very large and form additional subdomains; loops βB-βC (36 residues), βE-βF (75 residues) and βG-βH (223 residues) face the outer surface of the capsid (Fig. 1C, bottom). N- and C-terminal ends face the interior and the outer surface of the capsid, respectively. Fine structural details of capsid salient features, like spikes around the 3-fold axes (S3 regions), channels (pores) at the 5-fold axes (S5 regions), and distinct depressions at the 2-fold axes (S2 regions), could also be discerned (Fig. 1B).

#### A residue-by-residue comparison between cryo-EM LR values and crystallographic B-factors for the MVM capsid

The cryo-EM structural model of the wt MVMp capsid was then compared with the equivalent X-ray structural model of the same capsid previously determined at 3.25 Å resolution. Taking into account all 549 superimposed CP residues, the root mean square deviation (rmsd) was 0.62 Å for equivalent Cα (1.07 Å for all atoms). A residue-by-residue comparison of the cryo-EM structure and the crystal structure of the wt MVMp capsid showed only a few Cα atoms with rmsd > 0.80 Å (Fig. S2). This comparative analysis proved that the equilibrium conformations of this virus capsid in a hydrated crystal or in vitrified water are virtually indistinguishable.

The quality of different regions of the wt MVMp capsid cryo-EM map was determined using MonoRes (Vilas et al., 2018), which provides an accurate determination of LR (“Local resolution of cryo-EM maps with MonoRes,” 2018). MonoRes was chosen instead of other approaches including Fourier-shell-correlation-based ones because it simplifies the uncertainty in the locality (window size) and provides a conservative estimation of local resolution avoiding an overestimation of the resolution values. A program, Resol2Bfplot, was developed to extract and assign LR values for every atom. Relative LR values were calculated by dividing the absolute LR values obtained for the Cα atoms in each individual residue by the global (averaged), FSC-derived resolution of the map; these values were then appended directly to the atom coordinates in the standard PDB file (provided as Data S1). Quantitative differences between relative LR values in the cryo-EM structure of the CP subunit in the wt MVMp capsid were color-coded and represented on the atomic model (Fig. 2A). Several regions (peptide segments) in each CP molecule (labeled 1-10 in Fig. 2A), showed a comparatively lower resolution (relative LR values > 1.33; warmer colors in Fig. 2A). These regions corresponded mainly to protein loops that are mostly located on the outer capsid surface. Regions with the highest resolution (relative LR values <1.33; deep blue shades in Fig. 2A) corresponded to the β-barrel CP core.

**Figure 2.**
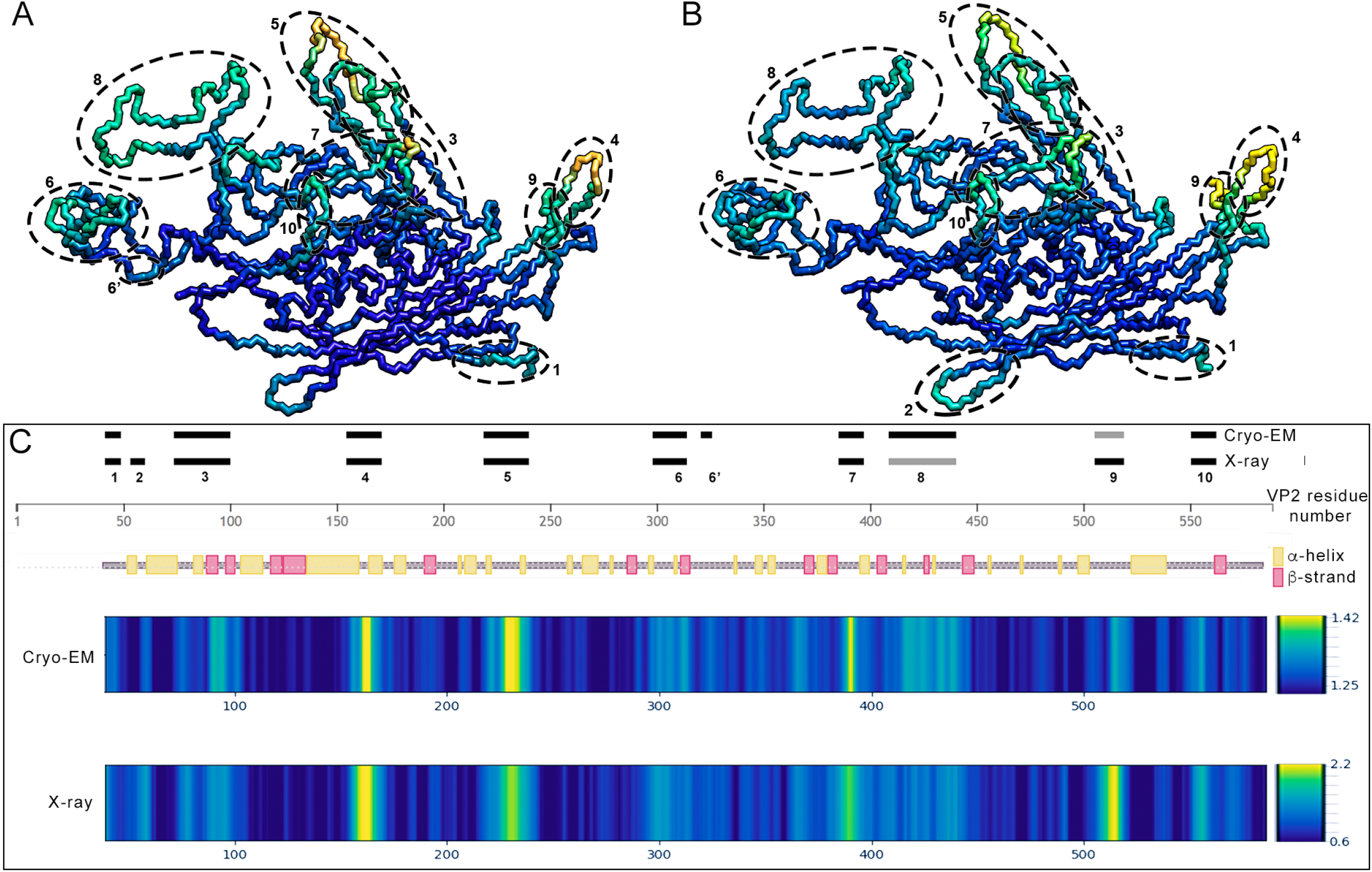
Comparison between cryo-EM LR values and X-ray crystallography B-factors for the wt MVM capsid. (A, B) Color-coded cryo-EM (A) and X-ray (B) structures of the wt MVM capsid according to relative LR values and relative B-factors per residue, respectively. Dashed ovals indicate peptide segments (1-10) with low LR values (A) or high B-factors (B). (C) Residue-by-residue comparison of cryo-EM relative LR values and crystallographic relative B-factors for the wt MVM capsid. Top: a grey line is used to indicate amino acid residue positions along the CP (VP2) sequence. α helices (yellow rectangles) and β strands (pink rectangles) are also indicated; black bars show the position of the segments (1-10) with low relative LR values in the cryo-EM structure and with high relative B-factors in the X-ray structure. Bottom: color distribution along the VP2 sequence is based on relative LR values (cryo-EM) or relative B factors (X-ray), as indicated in the color scales at right (values are given in Å or in Å^2^, respectively).

The relative LR values obtained by cryo-EM for each residue in the wt MVMp capsid were then compared with the relative B-factors previously obtained by X-ray crystallography for the same capsid (Kontou et al., 2005). For each residue, a relative B-factor was calculated by dividing the average B-factor for the main chain atoms of that residue by the average B-factor for all main chain atoms in the CP. Quantitative differences in relative B-factor were color-coded (with warmer colors representing higher B-values) and represented on the atomic structural model (Fig. 2B). A comparison between relative LR values (Fig. 2A) and relative B-factors (Fig. 2B) is shown in Fig. 2C. A clear match was found between the 10 solvent-exposed CP peptide segments that showed a comparatively lower resolution (relative LR values >1.33) in the cryo-EM structure (Fig. 2A, dashed ovals) and those with the highest B-factors (relative values >1.44) in the X-ray structure (Fig. 2B, dashed ovals). The same neat correspondence was found between regions with comparatively high resolution (low LR values) and low B-factors in the β-barrel CP core. This was not a merely qualitative coincidence. In a residue-by-residue, linear representation of LR values (cryo-EM) *versus* B-factor values (X-ray) along the CP sequence, a striking quantitative correlation was generally observed: compare the corresponding spectra of color hues in Fig. 2C (dark blue to deep yellow).

#### Comparison with HDX-MS results validates cryo-EM LR values and crystallographic B-factors as signatures of equilibrium dynamics for different structural elements in the MVM capsid in solution

HDX-MS is a gold standard method to investigate in solution the equilibrium dynamics of protein complexes including viruses (Bothner et al., 1998; Bothner & Hilmer, 2011; Bothner et al., 1999; Lewis et al., 1998; van de Waterbeemd et al., 2017; L. Wang & Smith, 2005; Worner et al., 2021). HDX-MS data are available for the wt MVMp capsid (van de Waterbeemd et al., 2017), which allowed to undertake a quantitative three-side comparison of the equilibrium dynamics of different regions and structural elements in the MVM capsid, as estimated by: i) the percent hydrogen-deuterium exchange for many individual peptide segments spanning the entire CP (VP2) sequence, determined by HDX-MS analysis of the MVM capsid in solution under defined conditions; ii) average relative LR values determined for the same peptide segments from cryo-EM data; and iii) average relative B-factors determined for the same peptide segments from X-ray crystallography data (Fig. 3). This comparison revealed an excellent match between CP peptide segments with a higher percent hydrogen-deuterium exchange (HDX-MS), those with a higher LR value (corresponding to lower resolution) (cryo-EM), and those with a higher B-factor (X-ray crystallography). The correlation between LR values and local differences in equilibrium dynamics in solution obtained by HDX-MS, a proven gold-standard technique for such analysis, validates the LR-based cryo-EM approach to investigate conformational fluctuations of biocomplexes at equilibrium.

**Figure 3.**
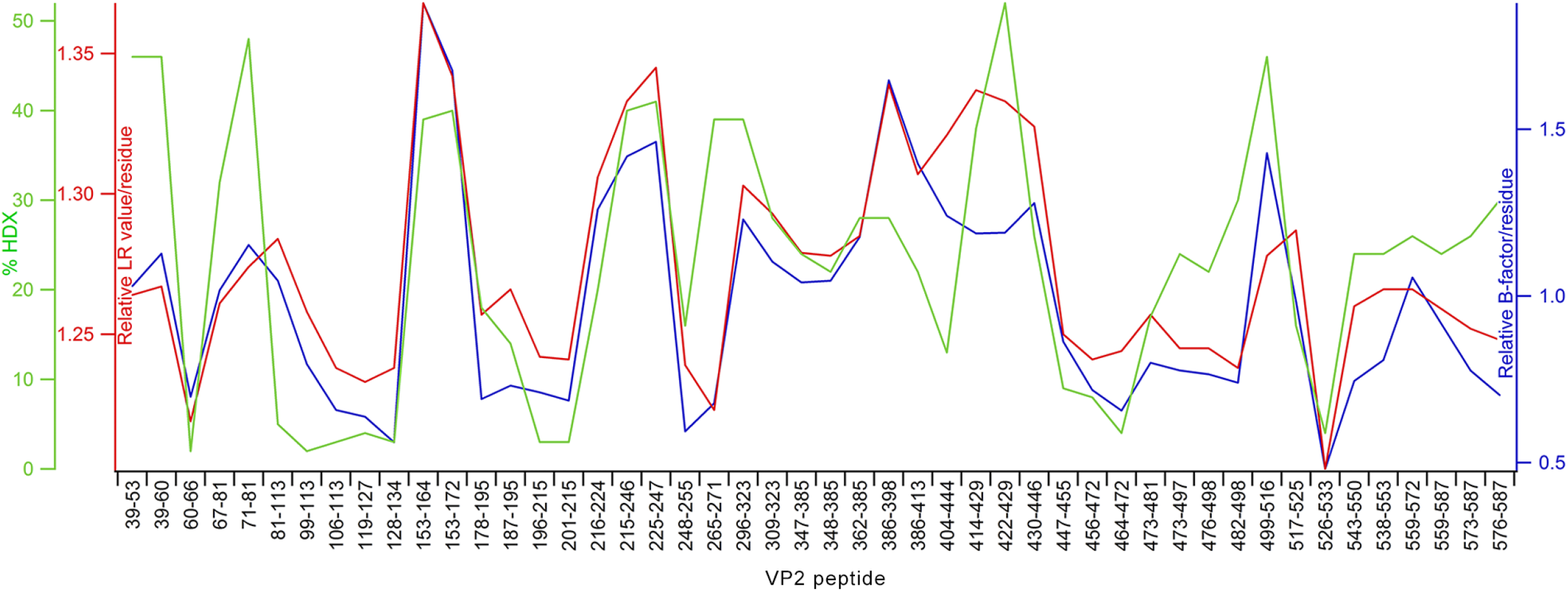
Superimposition of HDX-MS, cryo-EM and X-ray data for the wt MVM capsid. The Plot shows the percent hydrogen-deuterium exchange values (green), relative LR values (red) or relative B-factor values (blue) for the CP (VP2) peptide segments.

### A linkage between mutation-induced changes in mechanical elasticity and local changes in equilibrium dynamics of a virus capsid leading to impaired infectivity

The above validation of the LR-based cryo-EM approach to extract high-resolution information on the conformational fluctuations of single residues in a biocomplex should greatly facilitate structure-based mutational analyses to investigate the relationships between equilibrium dynamics and biological function, as proven next using MVM as a model system.

Using three different viruses (MVM, human rhinovirus and the human immunodeficiency virus), we had previously found evidence for an intimate relationship between mechanical stiffening and impaired conformational dynamics (transitions between equilibrium states and/or conformational fluctuations at equilibrium) (Carrillo et al., 2017; Castellanos, Carrillo, & Mateu, 2015; Castellanos et al., 2012; Dominguez-Zotes, Valbuena, & Mateu, 2022; Guerra et al., 2017; Valbuena & Mateu, 2015; Valbuena, Rodriguez-Huete, & Mateu, 2018). For MVM, different capsid-stiffening mutations impaired different biologically relevant transitions between conformational states (Castellanos et al., 2015; Castellanos et al., 2012), but it was unclear whether the equilibrium dynamics was also impaired. Two MVM capsids carrying single mutations (D263A or F55A) were selected for this study. These two amino acid substitutions are deleterious when individually introduced in the virus, virtually abolishing infection of host cells (Reguera, Carreira, Riolobos, Almendral, & Mateu, 2004). However, they are located far away from each other, and their biological effects are mediated by different mechanisms. Residue D263 (Fig. 4A, green) from five symmetry-related CPs delimits the base of the channel at each capsid five-fold axis. Mutation D263A impairs a conformational transition (Reguera et al., 2004) associated to externalization through the capsid channels of VP2 Nt segments carrying signals required for nuclear exit of MVM (Maroto, Valle, Saffrich, & Almendral, 2004; Valle, Riolobos, & Almendral, 2006). In contrast, residue F55 (Fig. 4A, blue) is located close to each capsid 2-fold axis at the center of each intertrimer interface, and mutation F55A hampers capsid assembly (Carrillo et al., 2017). Mechanical analysis using atomic force microscopy (AFM) revealed that both mutations resulted in capsid stiffening (Carrillo et al., 2017; Castellanos et al., 2012). We have now determined the cryo-EM structures of these two mutants to investigate, using the LR approach validated in the first part of this study, whether these deleterious, stiffening mutations impair conformational fluctuations of the MVM capsid at equilibrium.

**Figure 4.**
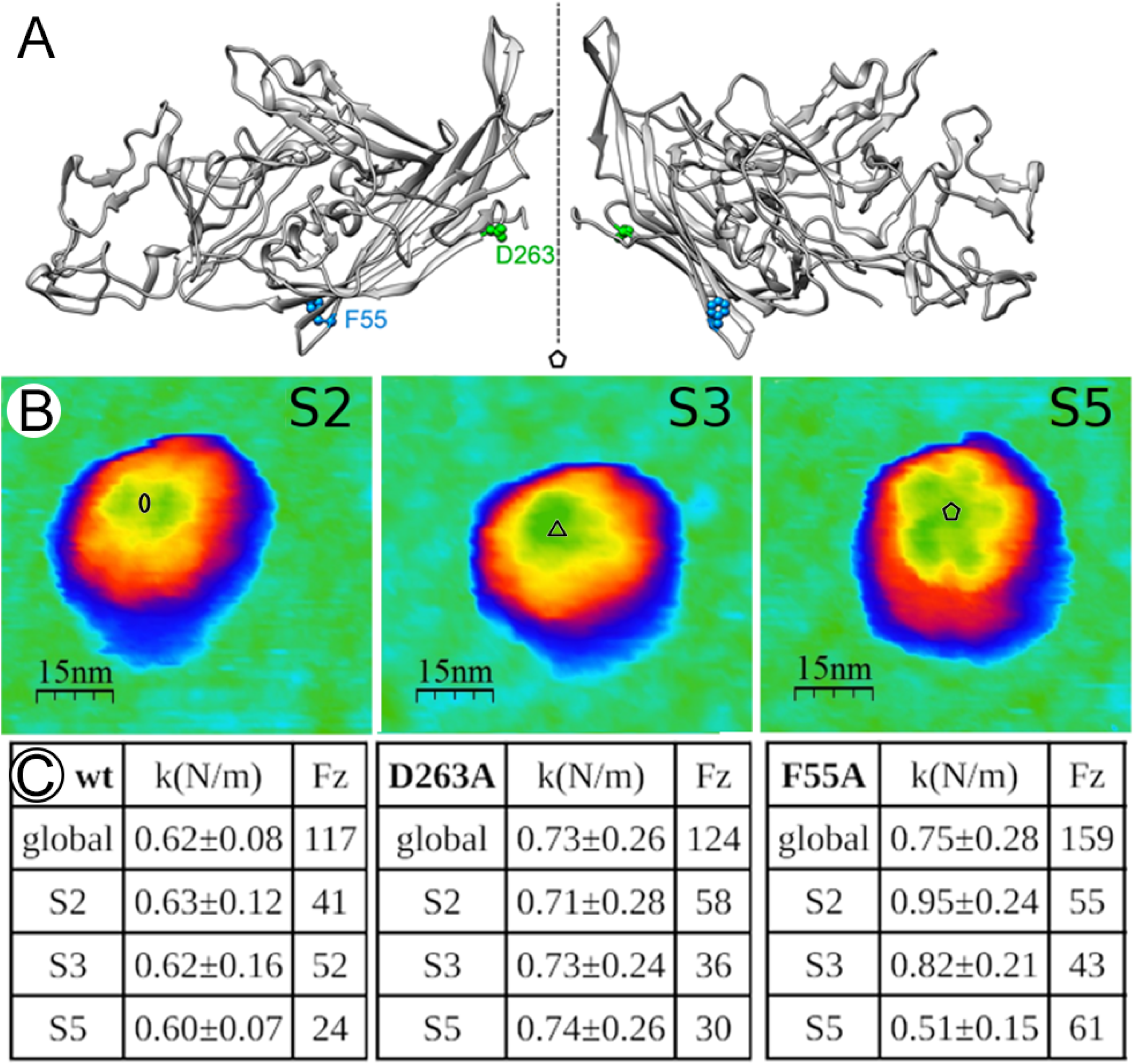
Mechanical stiffness of wt and D263A and F55A MVM capsids. (A) Side view of two CP subunits around a 5-fold axis (dashed line with a pentagon symbol) in the wt MVM capsid. Residues D263 (green) and F55 (blue) are indicated. (B) AFM images of individual D263A capsids with a 2-fold axis (S2 region) (left), 3-fold axis (S3 region) (middle) or 5-fold axis (S5 region) (right) on top. Oval, triangle and pentagon indicate approximate locations of 2-, 3- and 5-fold icosahedral symmetry axes, respectively. Scale bars are indicated. (C) Elastic constants (*k*, average value ± standard deviation) determined for wt, D263A and F55A capsids. Global, averaged *k* values without considering specific capsid regions; S2, S3 or S5, specific *k* values respectively determined for S2, S3 or S5 capsid regions. *k* values for wt and F55A have been previously published and are included here for completeness (Carrillo et al., 2017). *Fz*, number of capsid indentations used for analysis.

#### Effects of mutations on MVM capsid stiffness

Stiffening by the F55A mutation had previously been determined for VP2-only MVMp capsids, but stiffening by the D263A mutation had been determined for VP1/VP2 MVMp capsids, which differ from the former in the presence of disordered Nt extensions of a few (VP1) subunits. Although for the wt MVMp capsid and another mutant capsid (N170A) both VP2-only and VP1/VP2 versions showed the same stiffness (Guerra et al., 2017), a strict comparison of stiffness and dynamics between wt and mutants required the same capsid version to be used in all cases. Thus, the stiffness of the D263A VP2-only capsid was determined here as we had previously done for the wt and the F55A mutant VP2-only capsids.

High-resolution AFM imaging of individual D263A capsids from a highly purified preparation revealed the expected major topographic features. Different particles with a S2, S3 or S5 region on top were chosen for indentation (Fig. 4B), and force *versus* distance (*Fz*) curves were obtained. Shallow indentation of individual capsids elicited an elastic response. From the slope of the *Fz* curve, the elastic constant (*k*) was determined as described previously [(Castellanos et al., 2012; Ivanovska et al., 2004); see Methods]. The *k* values obtained for D263A capsids indented on a S2, S3 or S5 region were separately averaged for many particles, and taken as an indication of capsid stiffness when indented on that particular region (Fig. 4C).

The average (global) stiffness determined for the D263A capsid (disregarding the indented regions) was significantly higher than that of the wt capsid (Δ*k* = 18%), and similar to that previously obtained for the F55A capsid (Δ*k =* 21%). In both cases, capsid stiffening was anisotropic (Fig. 4C). D263A (located in S5 region, Fig. 4A) increased the stiffness of S5 regions (Δ*k =* 23%) more than that of S3 regions (Δ*k =* 18%) and S2 regions (Δ*k =* 13%). In contrast, F55A (located in S2 region, Fig. 4A) increased the stiffness of S2 regions (Δ*k =* 51%) more than that of S3 regions (Δ*k =* 32%) and did not increase the stiffness of S5 regions (Δ*k* = -15%).

#### Near-atomic-resolution cryo-EM structure of D263A and F55A mutant MVM capsids

Highly purified mutant capsid preparations were obtained, and structure determination was performed using exactly the same procedures followed for the wt capsid as described above. The D263A and F55A mutant MVM capsids were imaged by cryo-EM (Fig. 5A, B), and icosahedrally averaged maps were calculated at 3.36 Å resolution for D263A and at 3.26 Å for F55A, as estimated by the criterion of a 0.143 FSC coefficient (Fig. 5C, D; Fig. S1). The introduced amino acid substitutions D263A and F55A could actually be visualized in the cryo-EM density maps (especially for mutant F55A), which showed local differences in electron density consistent with each mutation (Fig. 5E, F). The cryo-EM atomic models of the F55A or D263A capsids were almost indistinguishable from that of the wt capsid, with respective rmsd of only 0.42 Å and 0.45 Å for Cα atoms (0.79 Å and 0.86 Å for all atoms). Thus, any difference in the capsid equilibrium conformation caused by either mutation (apart from the substituted residue itself) was exceedingly subtle.

**Figure 5.**
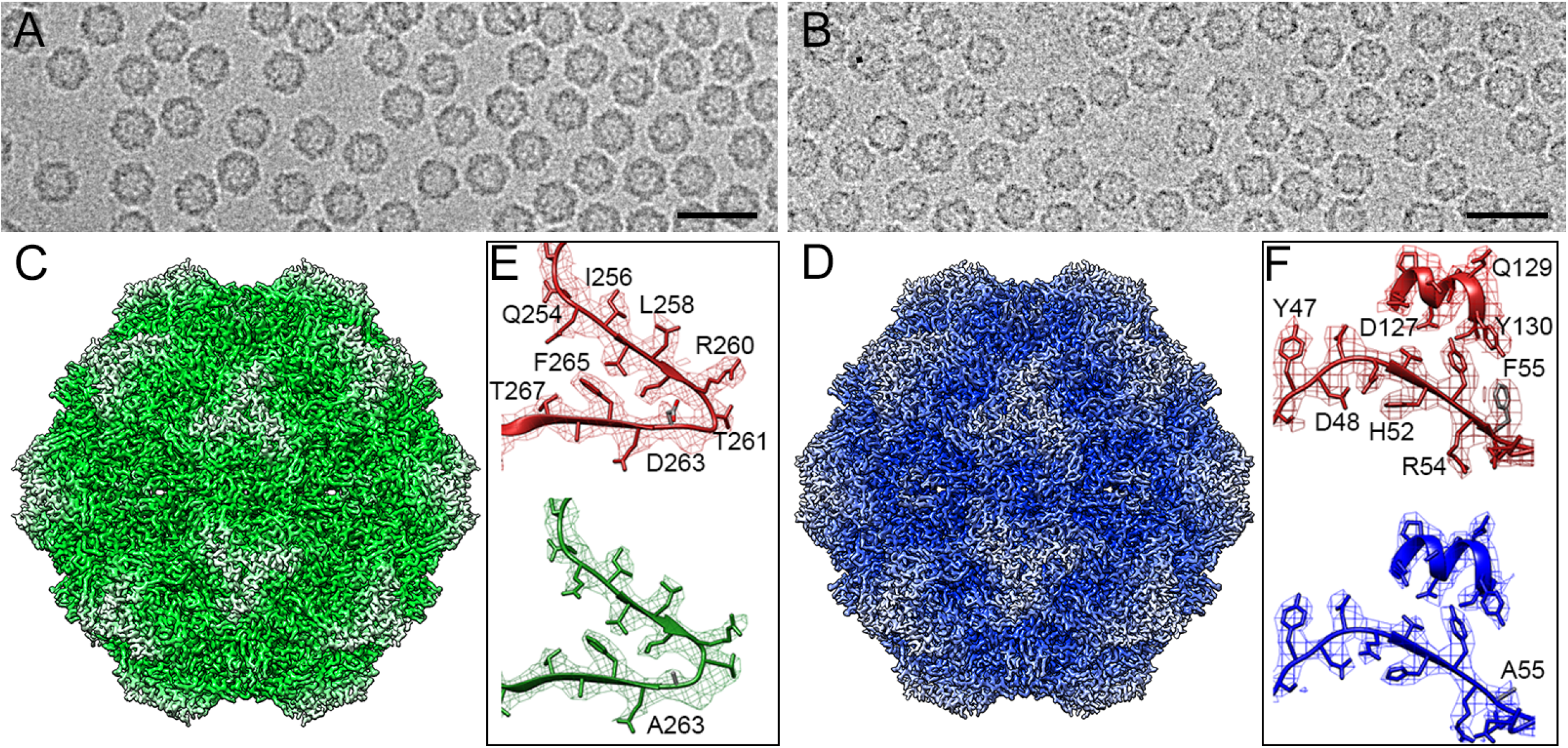
Cryo-EM structures of the D263A and F55A mutant MVM capsids. (A, B) Cryo-electron micrographs of D263A (A) and F55A (B) mutant capsids. Bar, 50 nm. (C, D) Radially color-coded atomic model of the D263A (C) and F55A (D) mutant capsids viewed along a 2-fold axis. (E, F) Cryo-EM densities for the capsid region around amino acid substitution D263A (E, green) or F55A (F, blue) compared to the corresponding region in the wt capsid (red).

#### Effect of D263A and F55A mutations in the equilibrium dynamics of the MVM capsid detected by cryo-EM

Despite the equilibrium conformations of wt, F55A and D263A MVM capsids were virtually indistinguishable, differences in local resolution in the corresponding cryo-EM density maps revealed many small, but significant, changes in capsid equilibrium dynamics (Fig. 6A). In general, conformationally more dynamic capsid regions in the wt capsid became less dynamic in the mutant capsids (Fig. 6A, B). Clear impairment of equilibrium dynamics was found around the mutated residues (positions 55 or 263) (Fig. 6B, red arrows) but also in many other capsid regions, some of them located very far from the mutated residue (Fig. 6B). Most of quenched dynamic regions involved capsid surface loops (Fig. 6B) distributed along the whole CP sequence (Fig. 6C).

**Figure 6.**
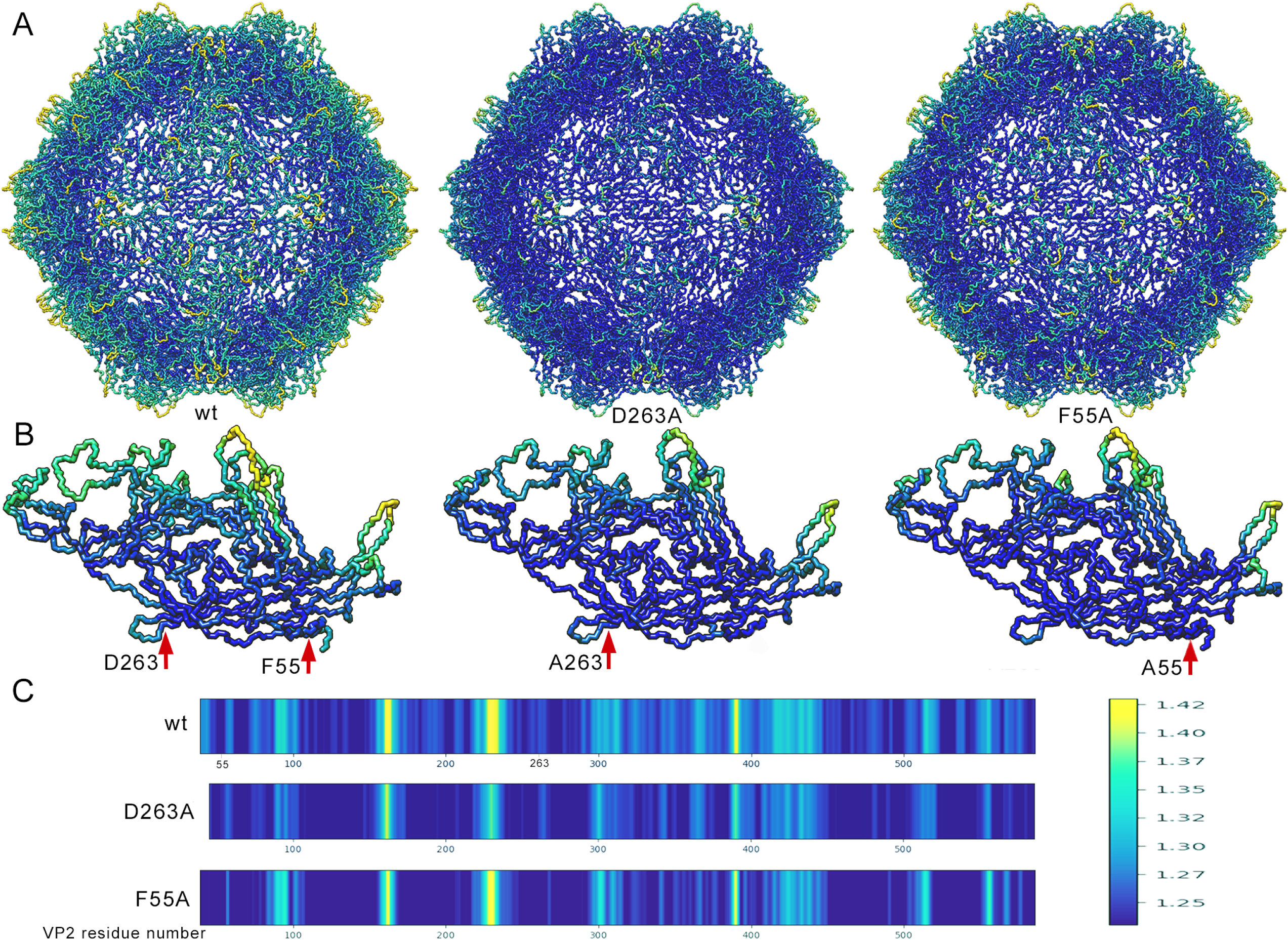
Effect of D263A and F55A mutations in the equilibrium dynamics of the MVM capsid as determined by cryo-EM. (A) Color-coded cryo-EM structures of wt (left), D263A (middle) or F55A (right) MVM capsids according to relative LR values per residue. (B) Cryo-EM structures of the wt (left), F55A (middle) or D263A (right) CP (VP2) subunit (color coded as in Figure 2A). Positions of wt residues D263 and F55 (left), or mutant residues A55 (middle) or A263 (right) are indicated. (C) Residue-by-residue comparison of the relative LR values for the wt, D263A and F55A MVM capsids. Color distribution along the VP2 sequence is based on relative LR values as indicated in the color scale at right (values are given in Å).

#### Comparative analysis of the effects of the D263A and F55A mutations on the mechanical stiffness and equilibrium dynamics of the MVM capsid

Comparison of the results obtained by AFM or cryo-EM for the wt, D263A and F55A capsids supports the hypothesis that an overall increase in capsid stiffness (Fig. 4) is intimately linked to an overall reduction in capsid equilibrium dynamics (Fig. 6). Moreover, when the effects of the D263A or F55A mutations on the equilibrium dynamics of specific regions were compared, some small differences could be detected. If confirmed by further analysis, some of those differences in local dynamics may underlie the different mechanisms by which those two mutations impair virus infectivity.

To sum up, the validated LR-based cryo-EM approach to obtain high-resolution information on the equilibrium dynamics of a viral capsid provided strong support to the proposal that the deleterious effect of capsid-stiffening mutations acting at different stages of the infectious cycle is ultimately related to their impairment of the conformational dynamics of the MVM capsid.

## Discussion

For a macromolecular complex, differences between atoms and structural elements in both relative B factors obtained by X-ray crystallography and relative LR values obtained by cryo-EM depend ideally on differences in their conformational dynamics. Both B factors and cryo-EM data have been used to extract dynamics information. However, crystallographic B-factors may be influenced by the general static disorder (slightly different relative orientations/positions of individual molecules in the crystal) and packing forces (which will impair the dynamics of regions involved in crystal intermolecular contacts). Likewise, LR values are influenced also by local damage during sample irradiation or inaccurate object orientation during image processing. In the face of these potential limitations, this study aimed to validate by referring to a gold-standard solution technique whether differences in relative LR values in cryo-EM maps could actually provide a reliable signature for small-scale differences in conformational (equilibrium) dynamics between different regions of a biomolecular complex. While both LR in cryo-EM and B-factors in X-ray crystallography are intrinsically dependent on dynamics, the factors that may affect one parameter or the other are different. Thus, the excellent correlation between cryo-EM relative LR values and crystallographic relative B-factors strongly support the view that both parameters are good indicators of local differences in conformational dynamics.

To discard the possibility that the MVM capsid might be an unusual complex for which those limiting factors had no significant effect on dynamics-related parameters, we tested the same approach for other viral and cellular protein complexes from the PDB (Berman et al., 2000) and EMDB (Lawson et al., 2016) repositories. Four different assemblies solved to atomic or near atomic resolution by both X-ray crystallography and cryo-EM were selected: β-galactosidase (Bartesaghi et al., 2018; Juers et al., 2000), dimethylformamidase [DMFase, (Arya et al., 2020; Vinothkumar, Arya, Ramanathan, & Subramanian, 2021)], apoferritin (Masuda, Goto, Yoshihara, & Mikami, 2010; Yip, Fischer, Paknia, Chari, & Stark, 2020) and porcine circovirus-2 [PCV-2, (Liu, Guo, Wang, Li, & Jiang, 2016; Mo et al., 2019)]. For every macromolecular complex a clear correlation between LR values and relative B-factors was found (Fig. 7).

**Figure 7.**
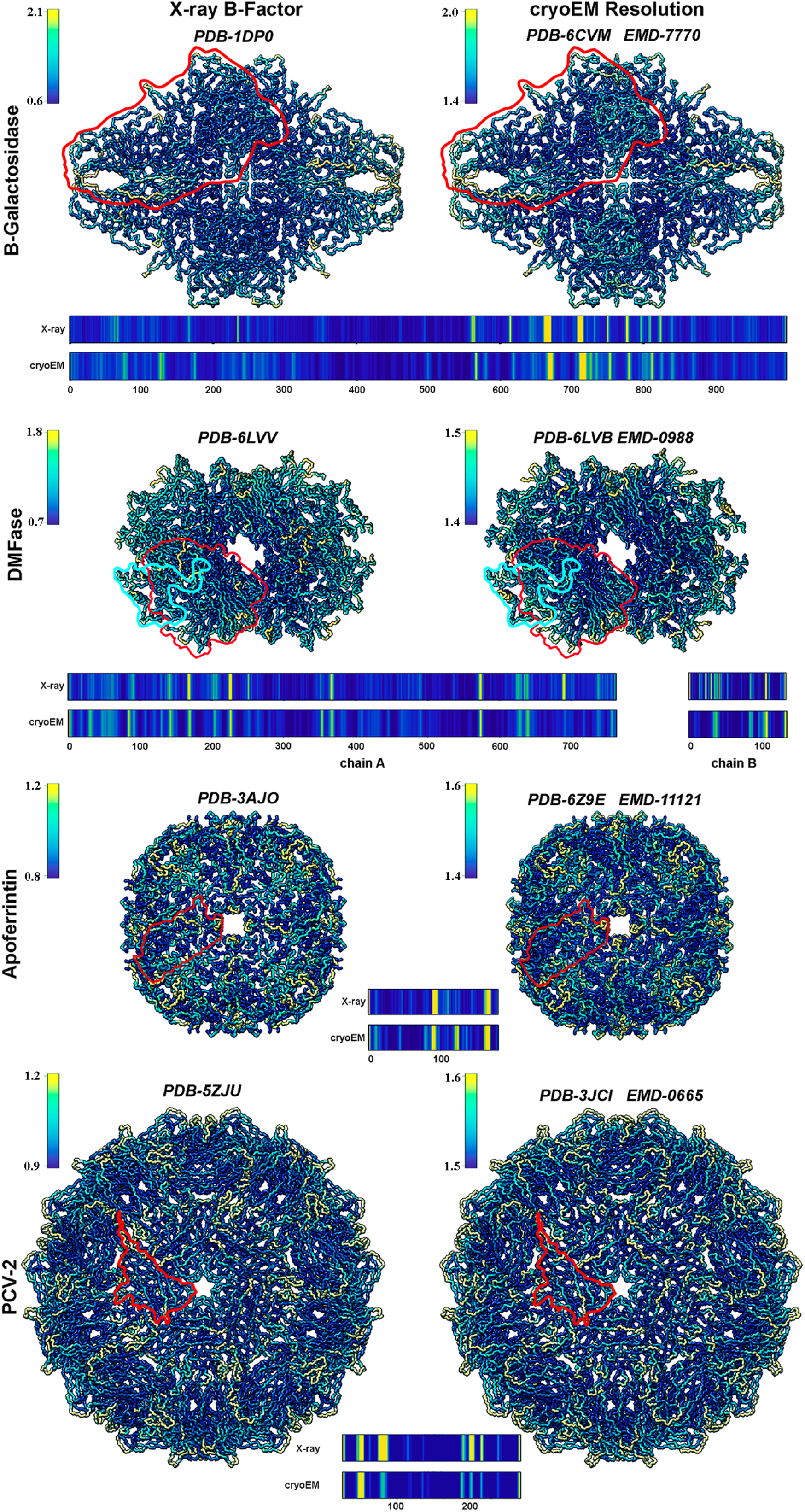
Comparison between cryo-EM LR values and X-ray crystallography B-factors for different protein complexes. X-ray (left) and cryo-EM (right) structures of (top to bottom) β-galactosidase, DMFase, apoferritin and PCV-2, color-coded according to relative LR values or relative B-factor values per residue, respectively. Bars below each pair of images represent a residue-by-residue comparison of the cryo-EM relative LR values and crystallographic relative B-factors along the protein sequence for the 4 complexes depicted. Color distribution along the protein sequence is based on normalized LR values or relative B-factors as indicated in the corresponding color scales.

More decisively, the comparison performed for the MVM capsid showed an excellent correlation between peptide-averaged LR values with the hydrogen-deuterium exchange percent values obtained in solution by HDX-MS (Fig. 3). This correlation indicates that differences in LR values can be directly related to differences in atomic fluctuations at equilibrium in solution.

Near-atomic-resolution determinations by cryo-EM are becoming a key to greatly expand and speed-up discovery of the structural basis for biological function of many viruses and cellular machines. Recently the cryo-EM structure of another parvovirus, AAV2, was determined at an unprecedented high resolution of 1.86 Å (Tan et al., 2018). The LR values ranged from 1.78 to 1.92 Å, but no dynamics information was extracted from those values. Comparing LR-based dynamic analysis of mutant and non-mutated forms of a biocomplex, especially using such highly detailed atomic structures, can now be used also to investigate in atomic detail structure-dynamics-related biological function relationships. Moreover, any remaining influence of technical limitations that could affect both the wt and the mutant will be removed when ΔLR = (LR_mutant_ - LR_wt_) values are obtained. As proof of concept, we applied here the cryo-EM LR approach, once validated in the first part of this study, to challenge a standing hypothesis for an inextricable relationship in viral particles between biologically relevant changes in mechanical stiffness and changes in equilibrium dynamics (discussed next).

We had previously found evidence for a relationship between global or local stiffening (determined by AFM) of virus particles and impaired conformational transitions and/or quenched equilibrium dynamics, and this linkage was rationalized in simple physical terms (Carrillo et al., 2017; Castellanos et al., 2015; Castellanos et al., 2012; Dominguez-Zotes et al., 2022; Guerra et al., 2017; Mateu, 2012; Valbuena & Mateu, 2015; Valbuena et al., 2018). A discriminating experiment using MVM capsid mutants was devised here to challenge the hypothesis of a linkage between mechanical stiffening and impaired equilibrium dynamics. In a previous study we had found many deleterious mutations in the MVM capsid that increased its stiffness (Carrillo et al., 2017; Castellanos et al., 2015; Castellanos et al., 2012). Some of those mutations (e.g., N170A, D263A) were located around the pores at the capsid 5-fold axes and exerted their deleterious effect by hampering a conformational transition associated to through-pore translocation of signal elements (Castellanos et al., 2012; van de Waterbeemd et al., 2017). In contrast, other stiffening mutations (e.g., F55A) far from the pores and close to the centers of the intertrimer interfaces at the capsid 2-fold axes had no effect on the pore-associated transition but impaired capsid assembly (Carrillo et al., 2017). We reasoned that, if an inextricable linkage between capsid stiffening and impaired equilibrium dynamics exists, every one of those capsid-stiffening mutations, irrespective of their differences in location or mechanism of action, should impair the capsid equilibrium dynamics.

In a seminal study of the nodavirus flock house virus (FHV), RNA-filled and empty capsids were shown to be structurally indistinguishable by X-ray crystallography; however, the RNA-filled capsid showed a markedly reduced conformational dynamics (Bothner et al., 1999). In the present study we found, using a cryo-EM-based LR approach, very similar results for two MVM capsid-stiffening mutations, D263A and F55A. Both mutations had exceedingly subtle effects on the capsid atomic structure but, based on lower LR values, they did substantially decrease the dynamics of many capsid elements. Moreover, the structural elements in which major reductions in dynamics were observed were very similar in the two mutant capsids. The observation that those capsid-stiffening deleterious mutations, irrespective of differences in location or mechanism of action, impair the dynamics of the capsid provides strong additional support for the hypothesis of an inextricable linkage between stiffness and equilibrium dynamics in a viral particle. Moreover, the high spatial resolution of the cryo-EM LR approach allowed the identification of differences on the effect of the tested mutations in the dynamics of specific loops and other capsid structural elements.

## Conclusions

Cryo-EM is being increasingly used to determine the atomic structure of viruses and biomolecular complexes. In the present study, direct comparison with HDX-MS results in solution validates LR values obtained from cryo-EM density maps as indicators of dynamic regions in biomolecular assemblies. The LR-based cryo-EM approach presents several strengths to investigate equilibrium dynamics:

i. It is relatively easy to implement on a general basis. For each new structure solved by cryo-EM at atomic or near-atomic resolution, we would like to propose that relative LR values are calculated and included in the atomic coordinate file deposited in the PDB.
ii. It provides near-atomic resolution information on differences in equilibrium dynamics between different structural elements, or after some structural modification is introduced (e.g., a mutation or bound ligand). In the latter case, the influence of technical artifacts will cancel out if ΔLR values are obtained.
iii. It complements other techniques to study equilibrium dynamics, including HDX-MS which may prove more accurate but is less resolutive. It may also be used to experimentally validate or guide even more detailed atomic resolution predictions by all-atom MD simulations.

This study provides also further support for an intimate linkage between changes in mechanical stiffness of a biomolecular complex and changes in its equilibrium dynamics. Information on equilibrium dynamics obtained by cryo-EM and/or AFM may help elucidate the physico-chemical mechanisms that underlie the biological function of viruses and cellular machines.

## Materials and Methods

### Recombinant plasmids, production and purification of MVM capsids

Recombinant plasmid pFB1-VP2 carrying the mutation D263A in VP2 of MVMp was used as a donor to construct the corresponding BM-VP2 bacmid using the baculovirus expression system (Invitrogen) (Reguera et al., 2004). BM-VP2 bacmids containing the wt VP2 gene or F55A or D263A mutant VP2 genes were used to produce the corresponding VP2-only MVMp capsids in H5 insect cells. Extensive purification of recombinant MVMp VP2-only capsids was performed essentially as previously described (Hernando et al., 2000). Purity and quality of the capsid preparations was assessed mainly by EM.

### Cryo-EM and data collection

Purified wt or mutant D263A or F55A MVMp capsid samples (5 μl) were applied onto R2/2 300 mesh copper grids (Quantifoil Micro Tools, Germany) and vitrified using a Leica EM CPC cryofixation unit. Data were collected on a FEI Talos Arctica electron microscope operated at 200 kV and images recorded on a FEI Falcon II detector for wt and D263A, and on a FEI Falcon III detector for F55A operating in linear mode. A total of 4,209 (wt), 3,002 (D263A) and 4,114 (F55A) movies were recorded at a calibrated magnification of X73,000 for wt, D263A and F55A, yielding in all cases a pixel size of 1.37 Å on the specimen. For wt and D263A, each movie comprises 23 frames with a dose rate of 1.59 e^-^ Å^-2^ per frame with an accumulated dose of 36.57 e^-^ Å^-2^; for F55A, each movie comprises 36 frames with a dose rate of 1 e^-^ Å^-2^ per frame with an accumulated dose of 36 e^-^ Å^-2^. Data acquisition was performed with EPU Automated Data Acquisition Software for Single Particle Analysis (ThermoFisher) with three shots per hole at -0.50 µm to -4.0 µm defocus for all samples.

### Image processing

All image-processing steps were performed using Scipion (de la Rosa-Trevin et al., 2016) package software. Movies were motion-corrected and dose-weighted with Motioncor2 (Zheng et al., 2017). Aligned, non-dose-weighted micrographs were then used to estimate the contrast transfer function (CTF) with CTFfind4 (Rohou & Grigorieff, 2015). All subsequent image processing steps were performed using RELION 2.1 (Scheres, 2012). 2D averages obtained from preliminary datasets were used as references to automatically pick the micrographs and a total of 997,001 (wt), 941,403 (D263A) and 1,015,264 (F55A) particles were extracted. 2D classification was performed for the three datasets and 841,632 (wt), 723,071 (D263A) and 914,218 (F55A) particles were selected to perform a 3D classification imposing icosahedral symmetry, using a model obtained from preliminary datasets low-pass filtered to 40 Å resolution as initial model. The best 210,854 (wt), 458,002 (D263A) and 171,062 (F55A) particles were included in 3D auto-refinement imposing icosahedral symmetry; yielding maps with an overall resolution at 3.42 Å (wt), 3.32 Å (D263A) and 3.22 Å (F55A) based on the gold-standard (FSC = 0.143) criterion. Local resolution was estimated using Xmipp3 MonoRes with two half volumes from the 3D auto-refinement analyzing the resolution range from the Nyquist limit of 2.74 Å to lower 10 Å limit resolution (Vilas et al., 2018).

### Model building and refinement

The structure of MVM [PDB ID: 1Z14; (Kontou et al., 2005)] was used as template to build a homology model. The model was first manually docked as a rigid body into the density and followed by real space fitting with the Fit in Map routine in UCSF Chimera (Pettersen et al., 2004). The model was then manually adjusted in Coot (Emsley & Cowtan, 2004) to optimize the fit to the density. Then, real space refinement was performed in Phenix (Adams et al., 2010) using global minimization, morphing, simulated annealing, local grid search, atomic displacement parameter (ADP), secondary structure restraints, non-crystallographic symmetry (NCS) restraints, side chain rotamer restraints, and Ramachandran restraints. Refinement statistics are listed in Table 1.

### Model validation and analysis

The quality of the atomic models, including basic protein geometry, Ramachandran plots, clash analysis, was assessed and validated with Coot, MolProbity (Chen et al., 2010) as implemented in Phenix, and with the Worldwide PDB (wwPDB) OneDep System (https://deposit-pdbe.wwpdb.org/deposition).

### Comparison of relative LR values obtained by cryo-EM with relative B-factors obtained by X-ray crystallography

To compare relative LR values with relative B-factor values, a common framework was necessary. A residue-by-residue comparison was chosen to avoid over-interpretation and facilitate comparison of the results. Moreover, the residues are a natural structural basis convenient for establishing comparisons when atomic models are involved.

#### Relative B-factor values obtained by X-ray crystallography

A relative B-factor value was obtained for each residue by calculating the average B-factor for its main chain atoms, and dividing this value by the average B-factor for all main chain atoms in the molecule. Main chain atoms only were considered for the calculations because they have a higher weight in defining capsid conformational fluctuations.

#### Relative LR values obtained from cryo-EM maps

The atomic model was converted into a binary mask with the shape of the protein. Thus, LR values around each Cα could be averaged to obtain a single value per residue. The mask was created by placing spheres with a radius R centered at the Cα atom positions. A value of 1 was assigned to those voxels inside the spheres of radius R, otherwise a value of 0 was set. Because the mask was created from the atomic model, the residue that corresponds to each sphere is known.

An adequate value to the sphere radius R was assigned as follows. For a same map different local resolution algorithms can provide different LR values (Vilas et al., 2020), mainly because of the different locality of the algorithms. The local resolution method used here was MonoRes (Vilas et al., 2018), which make use of a Riesz Transform to estimate the resolution. The unidimensional Riesz transform, known as Hilbert Transform as kernel 1/πx, when x = 3 pixels the kernel goes close to zero. Thus, a coarse locality of 3 pixels was chosen for using MonoRes. The sampling rate of the current detectors are around 1 Å/pixel or even lower. This pixel size is in the range of the locality of MonoRes, which justified the choice of a radius R = 3 Å for the spheres in the mask.

### Atomic force microscopy

AFM hardware and software were from Nanotec Electrónica. AFM imaging and quantification of mechanical stiffness were performed as previously described (Castellanos et al., 2012; Ivanovska et al., 2004). AFM images were obtained in Jumping Mode (Moreno-Herrero, Colchero, Gomez-Herrero, & Baro, 2004) using RC800PSA cantilevers (Olympus) with a nominal elastic constant of 0.1 N/m. The actual elastic constant *k*_*c*_ for each cantilever was determined before each experiment as described previously (Sader, Chon, & Mulvaney, 1999). AFM images were processed using the WSxM software (Horcas et al., 2007). The stiffness of different capsid regions was determined by indenting individual capsids with a 5-fold, 3-fold or 2-fold symmetry axis close to the top of the particle at the indentation point, as determined by previous high-resolution AFM imaging (see Results). To keep within the range leading to an elastic response and avoid particle damage, only measurements that involved an indentation depth between 0.5 nm and 2.0 nm were considered. The elastic constant *k* for the indented capsid region was determined assuming that capsid and cantilever behave as a system of two ideal spring in series (Ivanovska et al., 2004).

## Acknowledgments

We thank Rocío Arranz and Javier Chichón of the Cryo-EM CNB/CIB-CSIC facility (Madrid), in the context of the CRIOMECORR project (ESFRI-2019-01-CSIC-16), for help with cryo-EM data acquisition. This work was supported by grants from the Spanish Ministry of Science and Innovation (PID2020-113287RB-I00) and the Comunidad Autónoma de Madrid (P2018/NMT-4389) to JRC, and by a grant of the Spanish Ministry of Science and Innovation (RTI2018-096635-B-100) to MGM. MGM acknowledges also an institutional grant from Fundación Ramón Areces. AO-E acknowledges a Juan de la Cierva research contract (FJCI-2015-24086) from the Spanish Ministry of Economy and Competitivity. MGM is an associate member of the Centre for Biocomputation and Physics of Complex Systems. The Centro Nacional de Biotecnología is a Severo Ochoa Center of Excellence (MINECO award SEV 2017-0712). The funders had no role in the study design, data collection and interpretation, or the decision to submit the work for publication.

## Data availability

The atomic coordinates were deposited in the Protein Data Bank with codes 7Z5D, 7Z5E and 7Z5F, for MVM wt, D263A and F55A respectively. The cryoEM density maps were deposited in the EM Data Bank with codes EMD-14519, EMD-14520 and EMD-14521, for MVM wt, D263A and F55A respectively.

## Code availability

Resol2Bfplot has been made publicly available in Xmipp (de la Rosa-Trevin et al., 2013) and has been integrated in Scipion (de la Rosa-Trevin et al., 2016) (http://scipion.cnb.csic.es). A stand alone version can be found in (https://github.com/Vilax/bfactorLocalResolution).

---

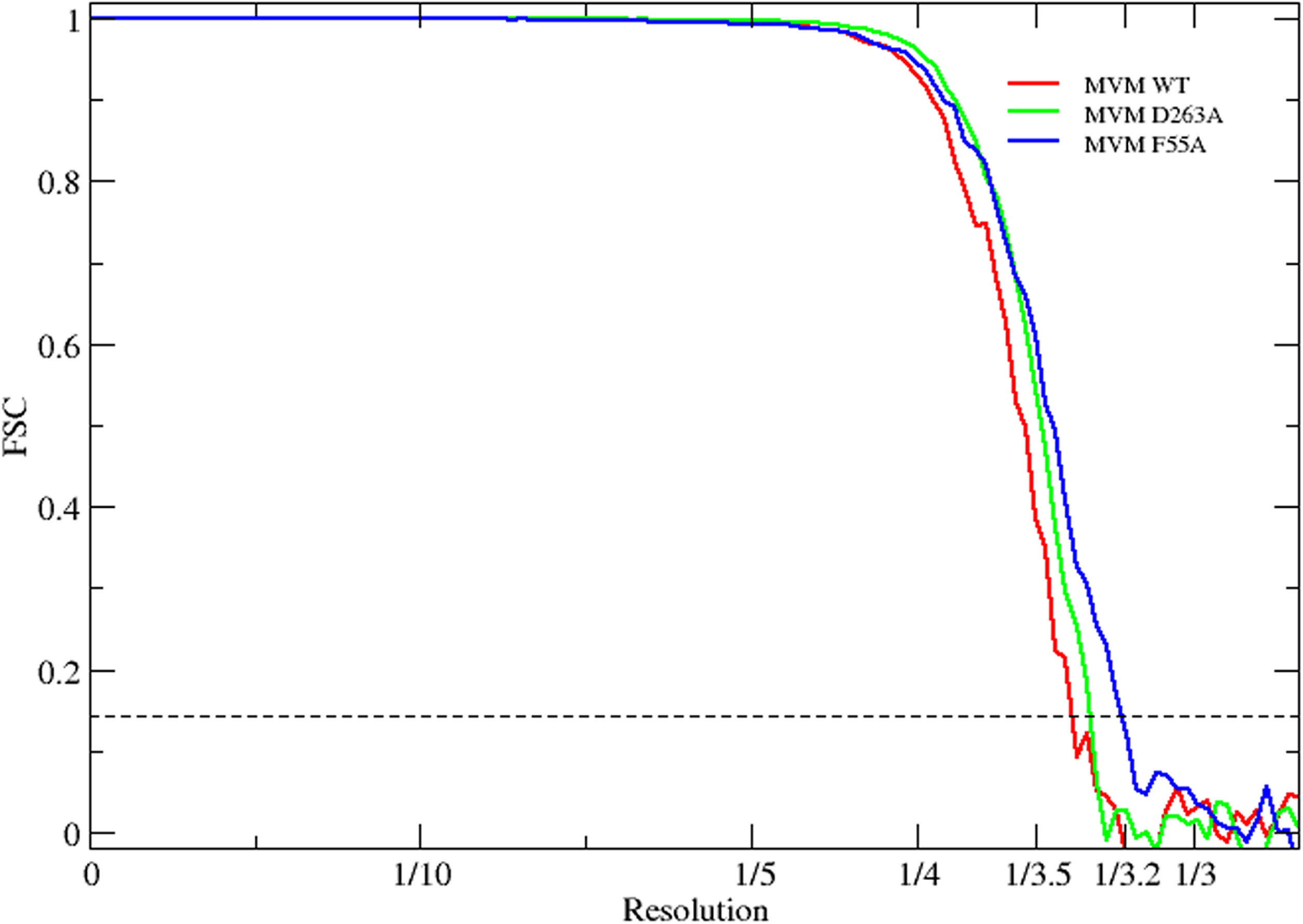

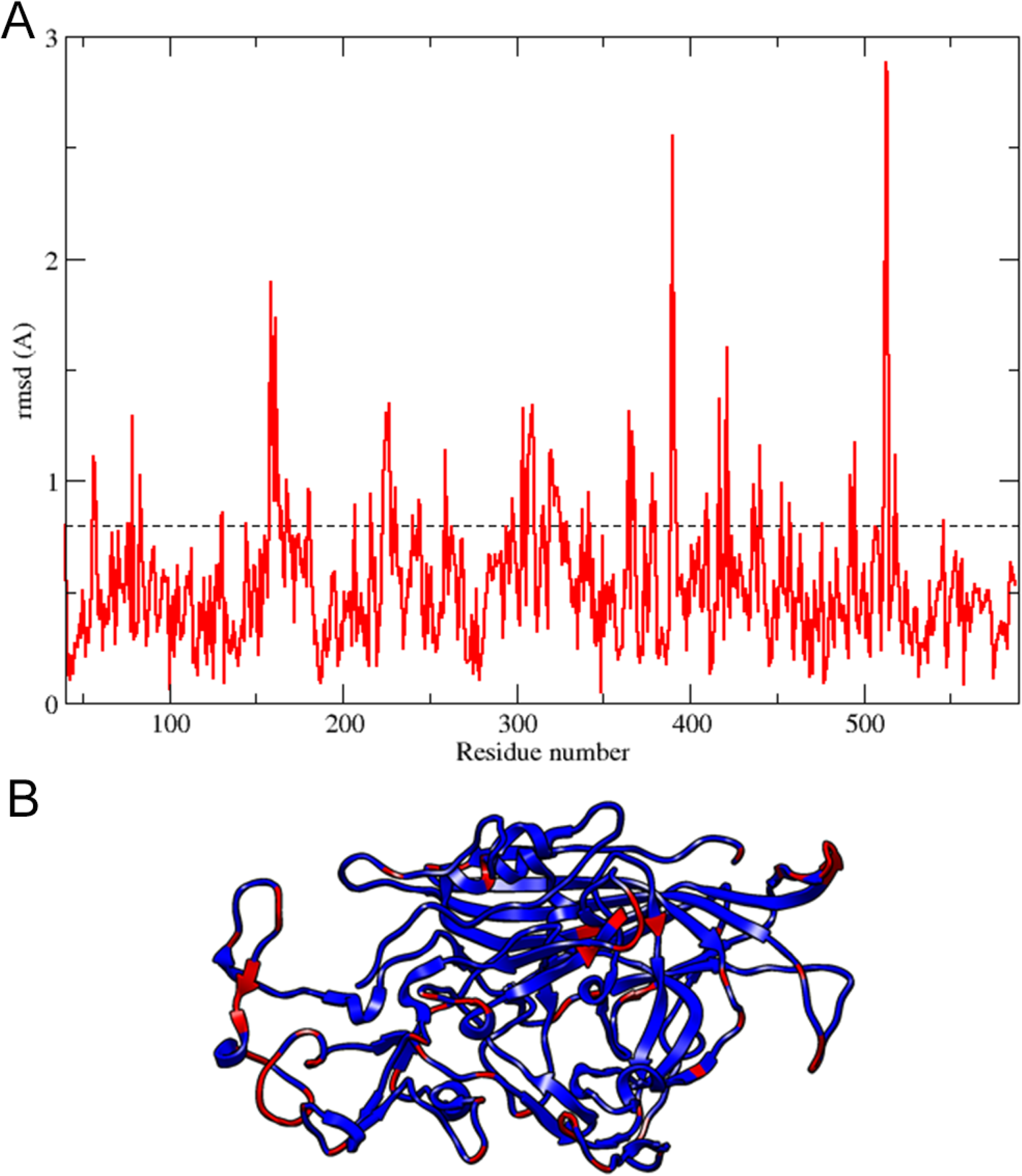

